# *Mimosa pudica*-derived zinc oxide nanoparticles preserve mesenchymal stromal cell viability, morphology, and osteogenic competence

**DOI:** 10.64898/2026.06.10.731413

**Authors:** AG Djuidje, P Belle Ebanda Kedi, AA Ntoumba, Marcus N. A. Fetzer, G Fonye Nuyfoni, G Chimi Tchoutchang, CC Nanga, UA Mintang Fongang, AK Tako Djimefo, BT Tabearuh Ayuk, MID Evouna, C Janiak, F Eya’ane Meva

**Affiliations:** Department of Pharmaceutical Sciences, Faculty of Medicine and Pharmaceutical Sciences, University of Douala, PO Box 2701, Douala, Cameroon; Department of Animal Biology and Physiology, Faculty of Sciences, University of Douala, PO Box 24157, Douala, Cameroon; Institute for Inorganic Chemistry and Structural Chemistry, Heinrich-Heine University, 40204 Düsseldorf, Düsseldorf, Germany; Department of Chemistry, Faculty of Sciences, University of Yaounde I, PO Box 812, Yaounde, Cameroon; Department of Chemistry, Faculty of Sciences, University of Douala, PO Box 24157, Douala, Cameroon

**Keywords:** *Mimosa pudica*, zinc oxide nanoparticles, mesenchymal stem cells, osteogenic differentiation, tissue engineering, regenerative medicine

## Abstract

**Introduction:** Musculoskeletal disorders remain a major cause of disability worldwide and require non invasive regenerative strategies that support tissue repair. Green-synthesized zinc oxide nanoparticles (ZnONPs) have attracted interest because of their biocompatibility and biological activity. This study investigated the synthesis of *Mimosa pudica*-derived ZnONPs (ZnO*MP*) and evaluated their effects on human bone marrow mesenchymal stromal cells (BM-MSCs).

**Methodology:** ZnO*MP* were synthesized using an aqueous extract of *Mimosa pudica* leaves and characterized by UV-Vis spectroscopy, FTIR spectroscopy, powder X-ray diffraction, SEM, EDS, and TEM. BM-MSCs isolated from human bone marrow were exposed to ZnO*MP*, plant extract, and synthesized ZnO nanoparticles. Cell metabolic activity was assessed by MTT assay after 1, 3, and 5 days. Cytoskeletal and nuclear morphology were analyzed by fluorescence microscopy and CellProfiler-based morphometry. Osteogenic differentiation was evaluated after 21 days using Alizarin Red S staining and quantification.

**Results:** Spectroscopic and microscopic analyses confirmed the successful formation of phytochemical-capped ZnO*MP* nanoparticles with nanoscale dimensions and specific elemental composition. ZnO*MP* maintained significantly higher metabolic activity than *Mimosa pudica* extract or ZnO at both 150 and 300 μg/mL. Morphometric profiling revealed that *Mimosa pudica* extract induced the most pronounced changes in nuclear morphology, reflecting enhanced nuclear plasticity and substantial remodeling of nuclear architecture, whereas ZnO*MP* preserved cellular and nuclear features closer to untreated controls. During osteogenic induction, ZnO*MP* did not impair matrix mineralization and preserved the ability of BM-MSCs to form a mineralized extracellular matrix.

**Conclusion:** *Mimosa pudica*-mediated ZnO nanoparticles combine favorable biocompatibility with preservation of mesenchymal stem cell morphology and osteogenic competence. These findings support their potential use as bioactive nanomaterials for musculoskeletal tissue engineering and regenerative medicine.

## Introduction

Musculoskeletal disorders (MSDs) comprise more than 150 conditions affecting bones, muscles, joints, and connective tissues, and represent one of the leading causes of disability worldwide (Cieza et al., 2021; Kiesel et al., 2024). MSDs are currently more common in high-sociodemographic index (SDI) regions, but their burden is gradually increasing in middle- and low-middle-SDI regions. (Zhou et al., 2024) These disorders include acute injuries such as fractures, sprains, and strains, as well as chronic and degenerative diseases including osteoarthritis, rheumatoid arthritis, osteoporosis, gout, and low back pain (Steinmetz et al., 2023; Zhou et al., 2024; Kiesel et al., 2024). MSDs are commonly characterized by pain, impaired mobility, reduced dexterity, and progressive loss of functional independence, thereby significantly affecting quality of life and work capacity (Zhou et al., 2024; Kiesel et al., 2024 and references therein). With the continuous increase in human life expectancy and the ageing of the global population, the prevalence of musculoskeletal diseases has risen markedly, creating a major challenge for healthcare systems worldwide (Nguyen et al., 2025; Yue et al.2025). In 2021, approximately 1.686 billion people were affected globally by MSDs, representing a 95% increase since 1990, and projections indicate that the number of cases may reach 2.161 billion by 2035 (GBD, 2017; IHME, 2026; Zhou et al., 2024). Similarly, osteoarthritis alone affected approximately 607 million people in 2021, with prevalence increasing sharply among individuals older than 55 years (Nguyen et al., 2025).

The burden of MSDs extends far beyond physical symptoms. These disorders are associated with chronic pain, functional disability, reduced mobility, psychological distress, and increased all-cause mortality. MSDs are currently recognized as the second leading cause of non-fatal disability worldwide and contribute substantially to years lived with disability (YLDs), with low back pain representing one of the most disabling conditions globally (Zhou et al., 2024; Zinabu et al., 2025). In addition to their effects on physical function, musculoskeletal diseases generate considerable socioeconomic consequences through healthcare expenditures, work incapacity, and long-term rehabilitation needs (Greggi et al.2024; Nguyen et al., 2025). Several factors contribute to the development and progression of MSDs, including ageing, obesity, reduced physical activity, depression, previous injuries, poor cardiovascular fitness, and occupational exposure to repetitive movements or prolonged awkward postures (Kiesel et al., 2024; Greggi et al.2024). Furthermore, studies suggest that women are more vulnerable to musculoskeletal disorders than men, likely because of hormonal influences and differences in pain perception and musculoskeletal physiology, while factors such as marital status, physical inactivity, unhealthy lifestyle behaviors, and depression also significantly contribute to the development of MSDs (Zinabu et al., 2025).

Despite the availability of pharmacological, non-pharmacological, and surgical interventions, the management of musculoskeletal diseases remains challenging. Conventional therapies such as analgesics, anti-inflammatory drugs, corticosteroids, bisphosphonates, and disease-modifying agents are often associated with limited long-term efficacy, adverse effects, drug interactions, and poor tolerance in older patients with multiple comorbidities (including diabetes, obesity, dementia, and stroke). (Kappenschneider et al., 2022; Tan et al., 2019; Rizzoli et al., 2011; Nguyen et al., 2025; Ziade et al., 2020). Senolytic agents such as dasatinib and quercetin, currently under investigation in preclinical models, have shown potential for selectively eliminating senescent cells and reducing pain-associated symptoms in osteoarthritis. (Gil et al., 2022; Nguyen et al., 2025) Romosozumab and denosumab have emerged as promising therapies for osteoporosis by not only increasing bone mineral density and reducing vertebral and non-vertebral fracture risk, but also improving muscle strength and lowering fall incidence, thereby highlighting the close relationship between bone and muscle health. (McClung et al., 2023; Rupp et al., 2022) Surgical approaches, including joint replacement and bone grafting, are generally reserved for severe cases but may be limited by frailty, postoperative complications, donor site morbidity, and incomplete tissue integration (Erezuma et al., 2021). Consequently, there is an urgent need for innovative and personalized therapeutic strategies capable of targeting the underlying pathological mechanisms while promoting tissue regeneration and functional recovery (Nguyen et al., 2025).

Recent advances in nanotechnology, regenerative medicine, and tissue engineering have opened promising perspectives for musculoskeletal disease management. Nanomaterials possess unique physicochemical and biological properties due to their nanoscale dimensions and high surface-to-volume ratio, enabling applications in targeted drug delivery, stem cell modulation, antimicrobial therapy, and tissue regeneration (Erezuma et al., 2021). Nanoparticles such as calcium phosphate and hydroxyapatite nanocrystals embedded within 1 nm triple-helix collagen fibrils can mimic the intricate nanoscale architecture of native bone tissue. (Lopes et al., 2018) Silver - Magnesium oxide nanocomposites mediated from *Talinum fruticosum* leaf extract revealed a marked improvement in antimicrobial activity compared to both the aqueous plant extract and monometallic silver nanoparticles (AgNPs). (Nyuyfoni Fonye et al., 2025) Gold nanoparticles (AuNPs) produced using *Eutrema japonicum* (Wasabi) possess antioxidant and anti-inflammatory activities. (Nanga et al., 2024) In particular, zinc oxide nanoparticles (ZnONPs) synthesized through green or biogenic approaches have attracted increasing attention because of their antioxidant, antimicrobial, anti-inflammatory, and anticancer properties (Berehu and Patnaik, 2024; Umar et al., 2025; Prashanth et al., 2024). Biogenic ZnONPs synthesized using plant extracts demonstrated significant biological activities, including radical scavenging effects, antibacterial activity against Gram-positive and Gram-negative bacteria, and concentration-dependent cytotoxicity against several cancer cell lines through reactive oxygen species (ROS)-mediated apoptosis pathways (Berehu and Patnaik, 2024). Importantly, green-synthesized nanoparticles have shown improved biocompatibility compared with chemically synthesized nanoparticles, highlighting their potential for safer biomedical applications (Maheswaran et al., 2024).

In the context of musculoskeletal tissue engineering, nanotechnology-based biomaterials and nanocomposites are increasingly being explored to improve bone regeneration and repair. Nanoclay-reinforced hydrogels and scaffolds have demonstrated promising osteogenic and angiogenic properties *in vivo*, supporting bone formation and tissue integration (Erezuma et al., 2021). Likewise, natural polymer-based nanocarriers and hybrid biomaterials combining organic and inorganic components have shown potential for improving drug delivery, osteogenesis, and cartilage repair while reducing inflammation and adverse effects (Zhou et al.2024). Recent studies also reported that green-synthesized ZnONPs nanoparticles derived from medicinal plants exhibit anti-inflammatory activity, while green Mg(OH)_2_ nanoparticles support mesenchymal stromal cell proliferation, and promote osteogenic differentiation, suggesting their relevance for bone regeneration and musculoskeletal tissue repair (Eya’ane Meva et al. 2025ab). Collectively, these advances indicate that nanotechnology and regenerative medicine may provide innovative and multifunctional therapeutic approaches capable of overcoming the limitations of conventional musculoskeletal disease treatments. This study describes the green synthesis and characterization of *Mimosa pudica*-derived zinc oxide nanoparticles and investigates their effects on the viability, morphology, and osteogenic competence of human bone marrow mesenchymal stromal cells, highlighting their potential for musculoskeletal tissue engineering and regenerative medicine

## Experimental

### Collection, authentication, and preparation of *Mimosa pudica* extract

Fresh leaves of *Mimosa pudica* L. were collected from the Botanical Garden of the Faculty of Medicine and Pharmaceutical Sciences, University of Douala, Cameroon. The plant material was authenticated at the Cameroon National Herbarium (Yaoundé, Cameroon), where a voucher specimen was deposited under accession number 54102/HNC. The collected leaves were thoroughly washed with running tap water followed by distilled water to remove dust and other surface contaminants. The cleaned leaves were then cut into small fragments (approximately 2 mm in length) to facilitate solvent penetration during extraction.

For aqueous extraction, 20 g of freshly prepared leaf material were immersed in 100 mL of distilled water preheated to 80 °C. The mixture was maintained under continuous magnetic stirring for 5 min using a thermostatically controlled hotplate. After cooling to room temperature, the extract was filtered through Whatman No. 1 filter paper and used for subsequent experiments.

### Green synthesis of ZnO nanoparticles (ZnO*MP*)

ZnO nanoparticles have been synthesized according to Eya’ane Meva et al., described protocols. (Eya’ane Meva et al., 2025a) briefly, a solution of 0.2 mol/L zinc nitrate hexahydrate (≥ 96%, CarlRoth) and a solution of 2 mol/L NaOH (≥ 97%, pellets, Merck) were prepared. 30 mL of *Mimosa pudica* aqueous extract was mixed with 70 mL zinc nitrate hexahydrate, and the solution was stirred for 10 min. The pH of the reaction mixture was adjusted by the dropwise addition of 2 mol/L NaOH at pH 12. Synthesis was monitored by a visual observation of the colloid suspension.

### Preparation of ZnO*MP* suspensions for biological assays

For biological investigations, purified ZnO*MP*s were dispersed in sterile distilled water and diluted to the desired concentrations prior to use. Suspensions were ultrasonicated to improve nanoparticle dispersion and minimize aggregation.

### Preparation of ZnO*MP* solutions for MTT assay

ZnO*MP* stock solutions (1000 μg/mL) were prepared in 99.9% ethanol according to a modified protocol adapted from Escheverry-Rendon et al. (Echeverry-Rendón et al., 2022) Before use, suspensions were ultrasonicated to ensure homogeneous dispersion. Working concentrations of 150 and 300 μg/mL were prepared and sterilized under ultraviolet irradiation for 30 min. These preparations were subsequently used for cytotoxicity and differentiation studies.

### Ultraviolet-visible spectroscopy (UV-Vis)

Optical characterization of the synthesized nanoparticles was performed using a P9 double-beam UV– visible spectrophotometer (VWR). Approximately 2 mL of nanoparticle suspension was scanned between 200 and 800 nm to identify characteristic ZnO absorption bands.

### Fourier transform infrared spectroscopy (FTIR)

FTIR analysis was carried out using a Bruker Tensor 37 spectrometer equipped with an attenuated total reflectance (ATR) accessory. Spectra were recorded in the 600–4000 cm□^1^ range to identify functional groups associated with phytochemical capping of the nanoparticles.

### Powder X-ray diffraction (PXRD)

The crystalline structure of the ZnO nanoparticles was analyzed using a Bruker D2 Phaser powder diffractometer equipped with Cu Kα radiation. Samples were prepared as thin films on low-background silicon sample holders prior to diffraction measurements.

### Scanning electron microscopy (SEM) and energy-dispersive X-ray spectroscopy (EDX)

Morphological examination of the nanoparticles was conducted using a JEOL JSM-6510LV scanning electron microscope operated at 20 kV. For elemental analysis, the instrument was coupled to a Bruker XFlash 410 silicon drift detector for EDX measurements. Prior to imaging, samples were sputter-coated with gold using a JEOL JFC-1200 fine coater.

### Transmission electron microscopy (TEM)

Transmission electron microscopy observations were obtained with a JEOL JEM-2100 microscope operating at 200 kV and equipped with a TVIPS F416 camera. Particle size measurements were performed using Fiji image analysis software.

### Isolation and expansion of bone marrow-derived mesenchymal stromal cells (BM-MSCs)

BM-MSCs were isolated from bone marrow obtained from two elderly female donors undergoing total hip replacement surgery, following informed consent and ethical approval procedures (ethical approval (187/18), University Hospital Würzburg, Germany). Mononuclear cells were separated by Ficoll density-gradient centrifugation according to Pereira et al. (Pereira et al., 2022)

### MTT cytotoxicity assay

Cell metabolic activity following ZnONP exposure was assessed using the MTT assay adapted from Pereira et al. (Pereira et al., 2022) BM-MSCs were seeded in 96-well plates at a density of 10 × 10^3^ cells/well and exposed to ZnO-NPs at concentrations of 150 and 300 μg/mL. Untreated cells, chemically synthesized ZnO, and plant extract alone served as controls. After incubation periods of 24, 72, and 120 h, culture media were removed and 10 μL of MTT solution (5 mg/mL) was added to each well. Following 3 h incubation at 37 °C, the generated formazan crystals were dissolved in 100 μL DMSO. Absorbance values were recorded at 540 nm using a microplate reader. Results were expressed as percentages relative to untreated control cells.

### Cell morphology assessment

Following 1 and 3 days of treatment with ZnONPs or controls, cells were washed with PBS and permeabilized with 0.1% Triton X-100. Blocking was performed using PBST containing BSA, after which cells were stained overnight with Phalloidin-iFluor 488 at 4 °C in darkness. Nuclear staining was achieved using DAPI. Fluorescence images were acquired using a Leica DMi8 fluorescence microscope. Automated quantification of cellular morphological parameters including area, compactness, aspect ratio, and solidity was performed using CellProfiler 4.2.5 as previously described with few modifications. (Pereira et al., 2022) Statistical analysis of morphological features was conducted using the Kruskal-Wallis test with significance set at p ≤ 0.05.

### Osteogenic differentiation assay

To evaluate the osteogenic potential of the synthesized ZnO*MP*, BM-MSCs were cultured in osteogenic differentiation medium for 21 days in the presence of nanoparticles and control treatments. Cells were seeded in 24-well plates at 2 × 10□ cells/cm^2^ until confluence was reached. Osteogenic differentiation medium consisted of DMEM supplemented with fetal calf serum, β-glycerophosphate, ascorbic acid-2-phosphate, and dexamethasone. Nanoparticles were added at a final concentration of 150 μg/mL and replenished during medium changes. After 21 days, cultures were fixed with 70% ethanol and stained with 2% Alizarin Red solution to visualize mineralized matrix deposition. Bright-field micrographs were recorded using a Leica DMi8 microscope. Quantification of incorporated dye was performed after extraction with cetylpyridinium chloride solution and absorbance measurement at 570 nm (Eya’ane Meva et al., 2025b).

### Statistical analysis

Experimental data were analyzed using GraphPad Prism software (version 9.1). Quantitative results were expressed as mean ± standard error of the mean (SEM). Statistical comparisons between groups were performed using two-way ANOVA followed by Tukey’s post hoc test, with p < 0.05 considered statistically significant.

## Results and discussion

ZnO*MP* nanoparticles were successfully synthesized through thermal activation of zinc ions and plant metabolites, followed by OH□-induced rapid nucleation of a characteristic white ZnO precipitate with plasmon resonance band ∼400 nm (Figure 1) that aligns with established metal-oxide synthetic precedents. (Eya’ane Meva et al., 2025a). Fourier transform infrared spectroscopy of isolated powders, demonstrated that the aqueous extract of *Mimosa pudica* and the biosynthesized ZnO*MP* nanoparticles (Figure 2) shared several absorption bands, indicating the involvement of plant-derived phytochemicals during nanoparticles formation. The broad band observed around 3290 cm□^1^ was attributed to O-H stretching vibrations of hydroxyl groups associated with phenolic compounds, flavonoids, and adsorbed water molecules. Absorption peaks detected near 2936 and 2850 cm□^1^ in both samples are attributed to asymmetric and symmetric C–H stretching vibrations of aliphatic groups, indicating the presence of organic constituents derived from the plant extract. The persistence of these signals after nanoparticle formation confirms that phytochemicals remained adsorbed on the ZnO surface as capping agents. The intense bands observed around 1642 cm□^1^ and 1537 cm□^1^ were assigned to carbonyl, amide, aromatic C=C, and N–H vibrations. The band at 1029 cm□^1^ were attributed to C–O and C–O–C stretching vibrations of alcohols, polysaccharides, or flavonoid structures. The slight variations in peak position and intensity after nanoparticle formation suggest interactions between zinc oxide and oxygenated phytoconstituents. Finally, the lower-frequency region below 700 cm□^1^, are generally characteristic of Zn–O stretching vibrations. Overall, the presence of similar vibrations in the spectra of the plant extract and ZnO*MP* nanoparticles demonstrate that bioactive compounds from *Mimosa pudica* played an important role as reducing, capping, and stabilizing agents during the green synthesis process (Maheswaran et al., 2024). Powder X-ray diffraction analysis (Figure 3) revealed broadened diffraction peaks indicating the presence of amorphous material when combining the *Mimosa pudica* extract and zinc nitrate.

**Figure 1.**
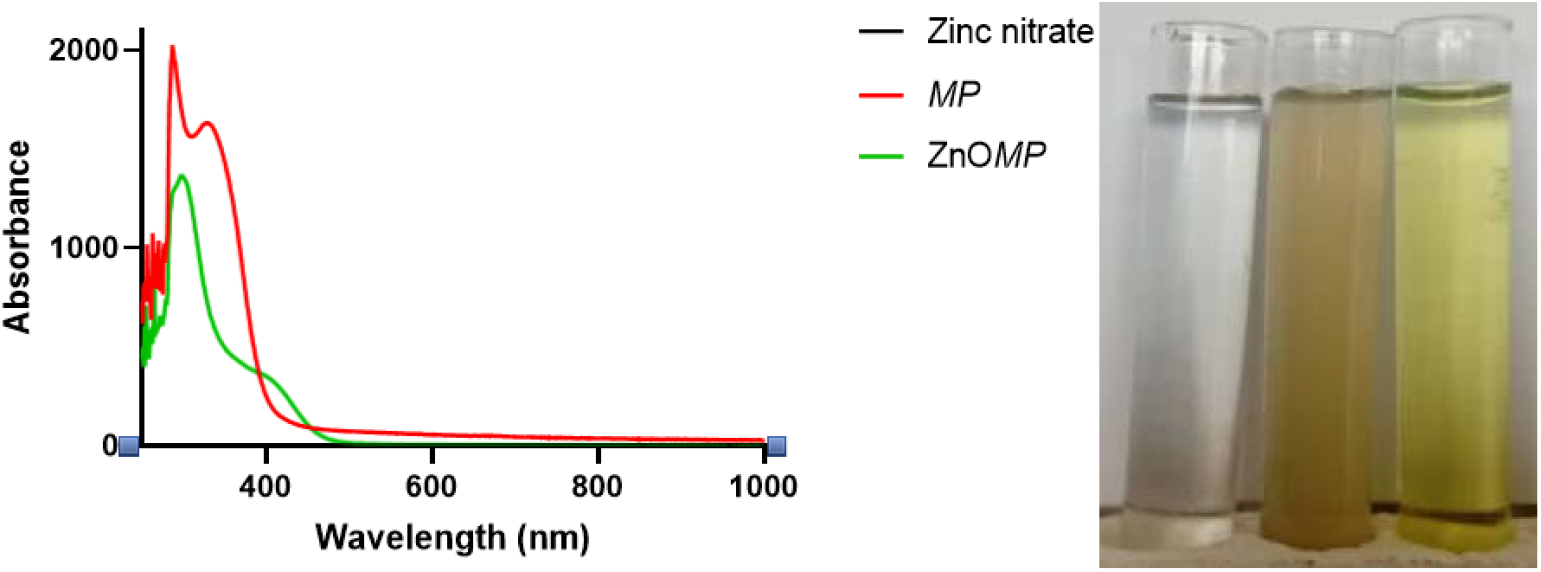
UV-Visible spectra (left) and visual observation of the colorations during the nanoparticle’s formation (right). Left: zinc nitrate, middle: *Mimosa pudica* plant extract, right: ZnO*MP*

**Figure 2.**
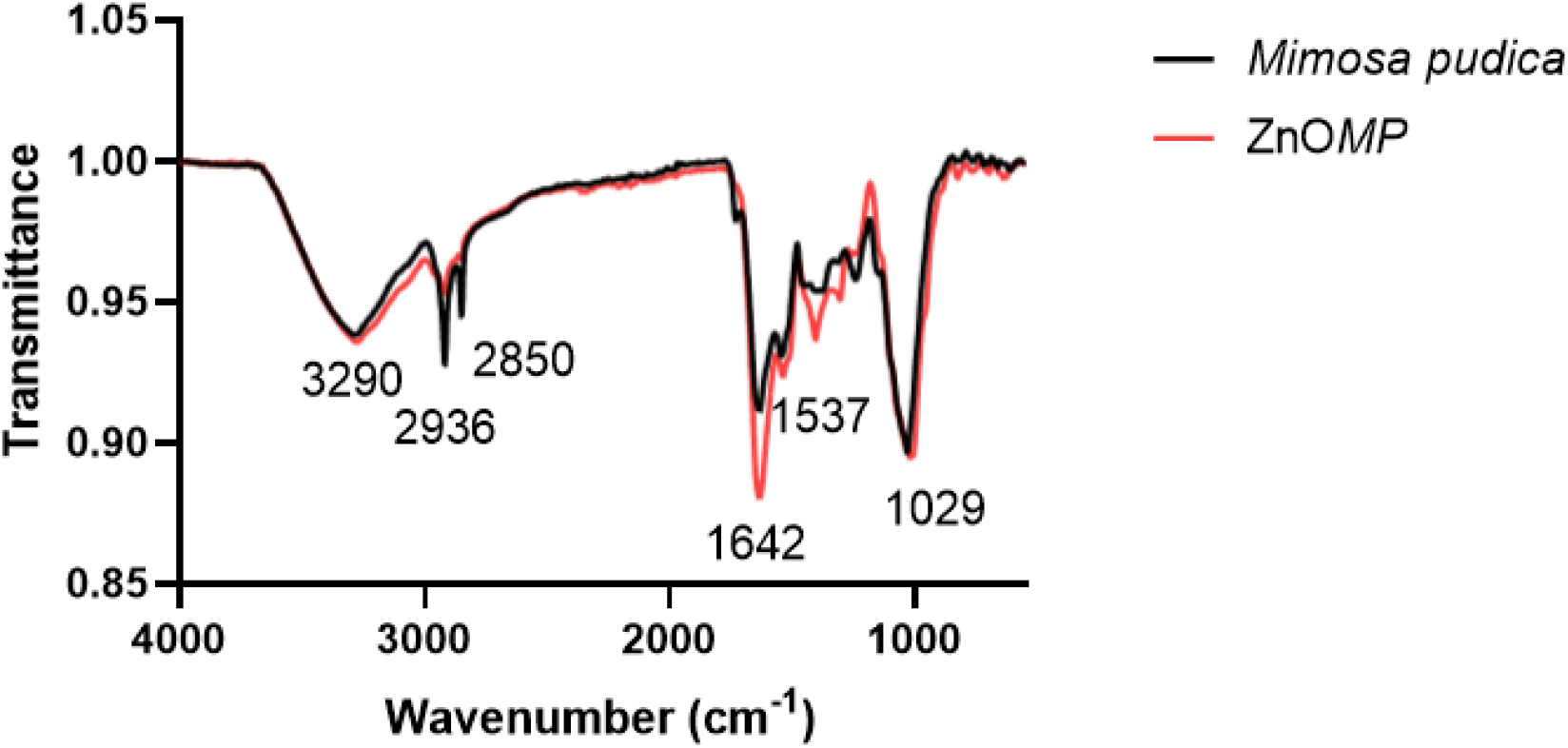
Fourier Transform Infrared Spectroscopy of *Mimosa pudica* and ZnO*MP*

**Figure 3.**
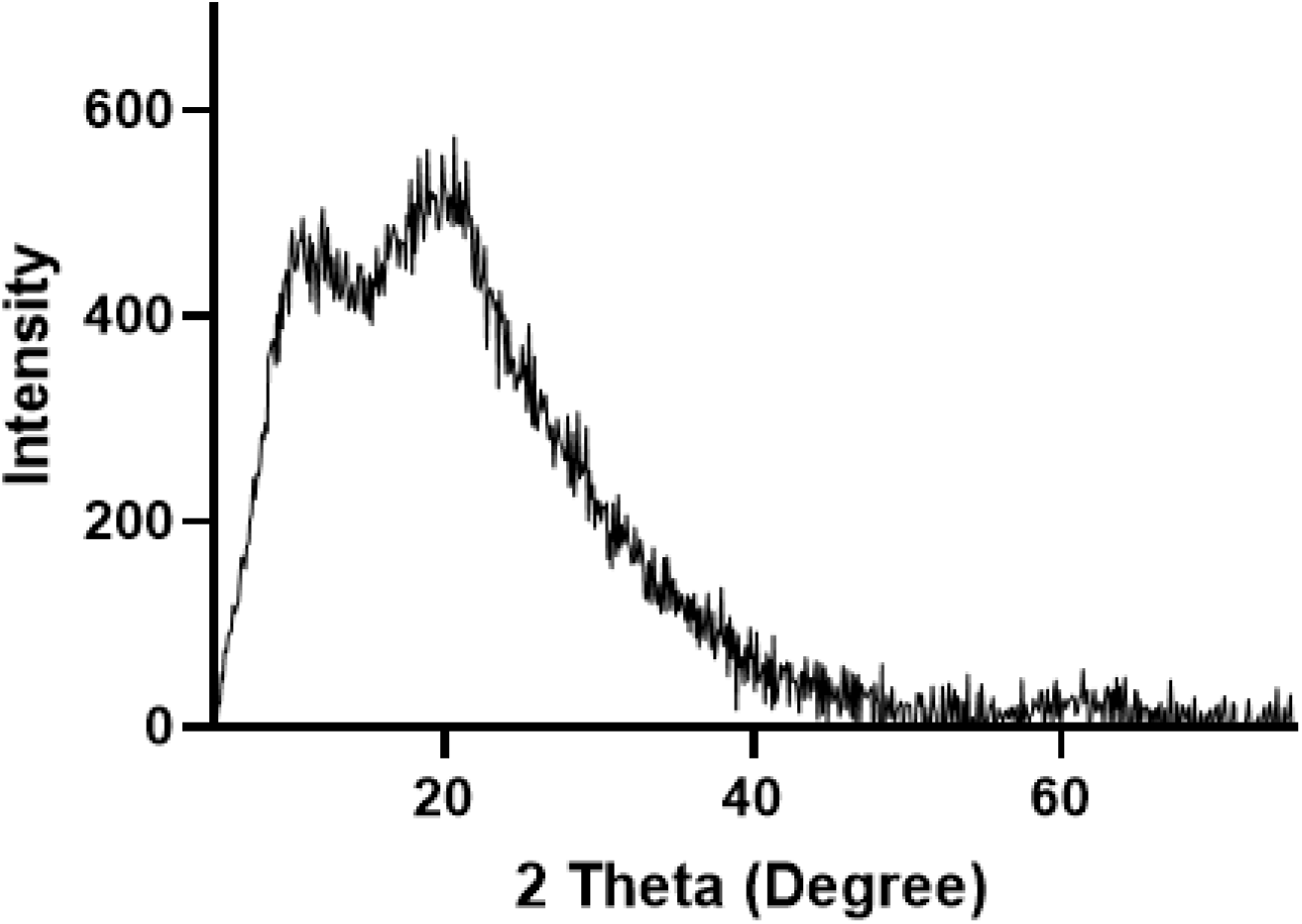
Powder X-ray Diffraction of ZnO*MP*

The scanning electron microscopy of the biosynthesized ZnO nanoparticles (Figure 4) revealed the formation of irregularly shaped aggregates composed mainly of plate-like and flake-like structures with heterogeneous particle dimensions. The particles appeared densely packed with rough and non-uniform surfaces, suggesting agglomeration during the drying process. Similar irregular and aggregated morphologies have frequently been reported for phytofabricated ZnO nanoparticles synthesized using plant extracts (Eya’ane meva et al., 2025a). The EDX spectrum confirmed the elemental composition of the synthesized material. The intense peaks corresponding to zinc (Zn) and oxygen (O) verified the formation of ZnO nanoparticles. The presence of carbon (C) as the dominant element (72.36 At%) is associated with organic compounds derived from the *Mimosa pudica* extract that remained adsorbed on the nanoparticle surface after synthesis. Minor phosphorus (P) signals detected in the spectrum may originate from naturally occurring biomolecules or mineral constituents present in the plant extract. The SEM and EDX analyses demonstrate that the green synthesis route successfully generated ZnO particles coated with phytochemical constituents, which may contribute to enhanced biological activity, colloidal stability, and biocompatibility. Transmission electron microscopy (Figure 5) confirmed the nanoscale nature of the synthesized material. Fine nanoparticles with dimensions of only a few nanometers were observed, comparable to previously reported phytomediated ZnO nanoparticles synthesized using *Aframomum citratum* extracts (Eya’ane Meva et al., 2025a).

**Figure 4.**
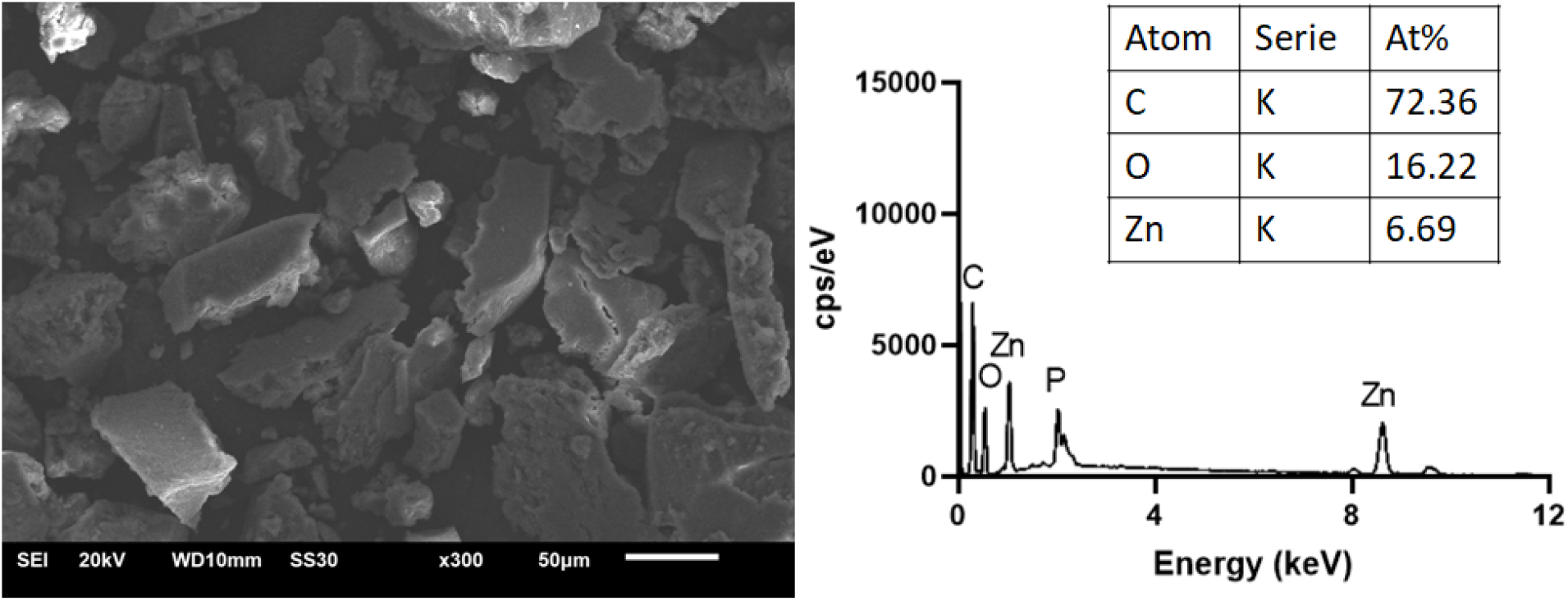
Scanning Electron Microscopy (SEM) and Energy Dispersive X-ray Spectroscopy EDX of ZnO*MP*

**Figure 5.**
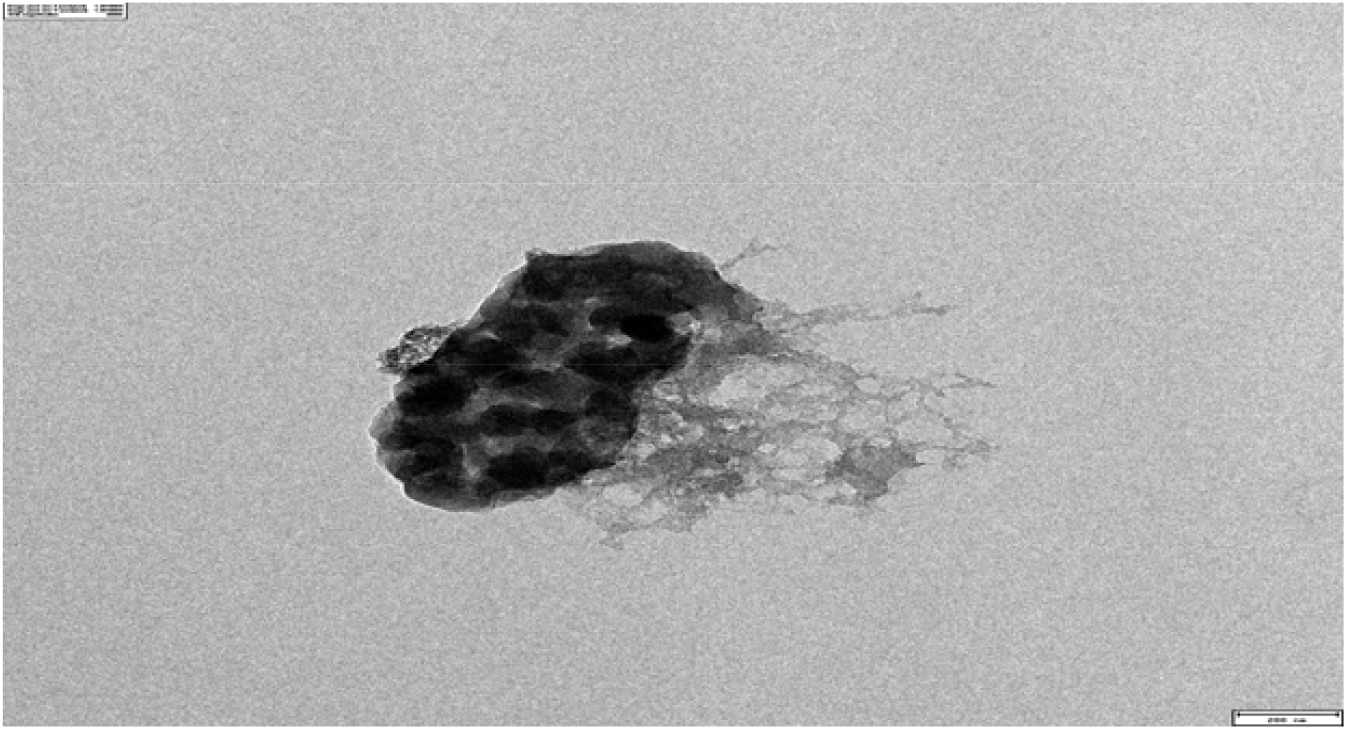
Transmission Electron Microscopy (TEM) of ZnO*MP*

The biological effects of ZnO*MP* were investigated through complementary analyses of metabolic activity, cellular morphology, nuclear architecture, and osteogenic differentiation. Together, these approaches provide a comprehensive assessment of both the immediate cellular response and the preservation of long-term mesenchymal stromal cell functionality. The MTT assay was used as an initial indicator of cellular metabolic activity and viability (Figure 6). Exposure to *Mimosa pudica* extract and chemically synthesized ZnO nanoparticles reduced cellular metabolic activity, whereas ZnO*MP* consistently maintained higher viability and promoted MSC proliferation. Similar improvements in cellular compatibility have previously been reported for phytochemical-coated magnesium hydroxide nanoparticles, suggesting that plant-derived surface molecules can substantially influence nanoparticle–cell interactions (Eya’ane Meva et al., 2025b).

**Figure 6.**
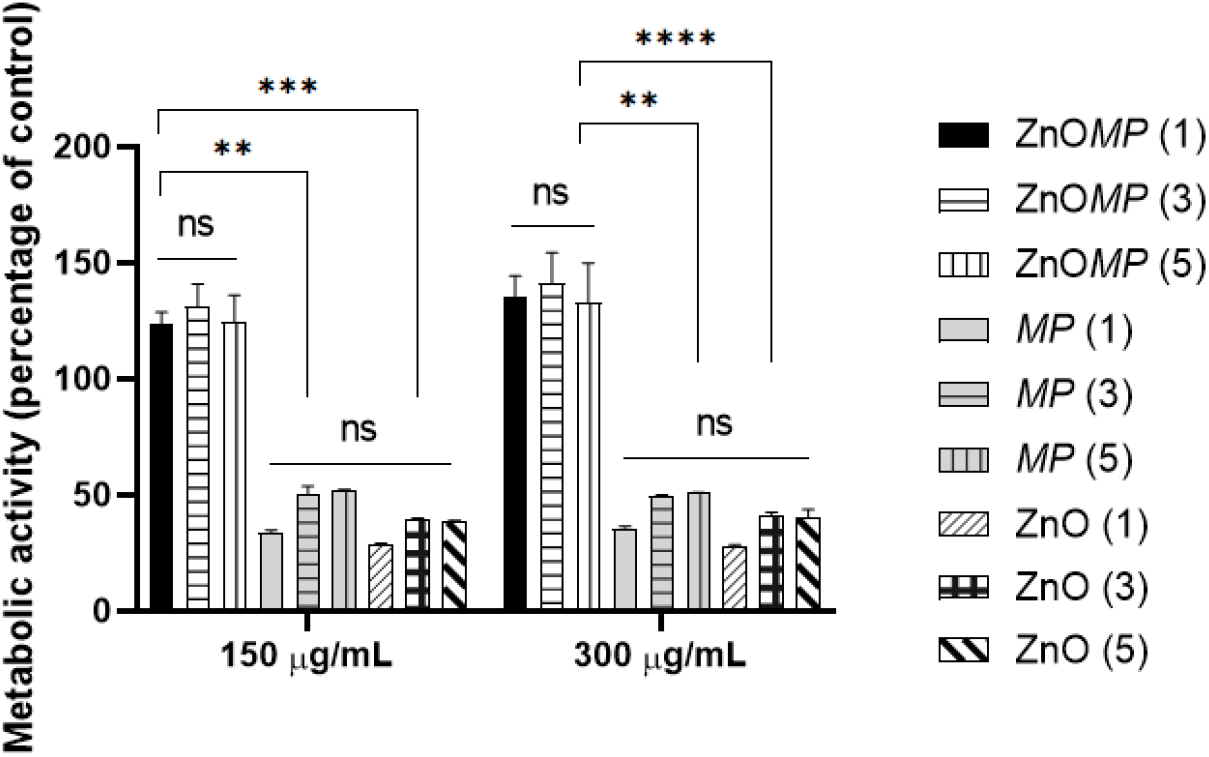
Cell viability / proliferation (MTT assay) of MSCs cultured under basal conditions and exposed to ZnO*MP, MP* and ZnO nanoparticles at 150 μg/mL and 300 μg/mL for 1, 3, and 5 days.

Morphometric of fluorescence microscopy images (Figure 7a) profiling further revealed that treatment-induced effects extended beyond metabolic activity and involved substantial remodeling of cellular architecture. Cell morphology is strongly regulated by cytoskeletal organization and mechanotransduction pathways. Changes in cell spreading, adhesion, and shape therefore provide sensitive indicators of cellular stress and functional adaptation (Berdiaki et al., 2024). Cell area, perimeter, maximum radius, and Feret diameters (Figure 7b) collectively describe the extent of cell spreading and adhesion to the substrate. Under normal culture conditions, MSCs adopt a fibroblast-like morphology characterized by extensive actin stress fibers and broad cell–substrate contacts. Because MSC differentiation and matrix production are strongly dependent on cytoskeletal tension and adhesion-mediated signaling, reductions in these descriptors generally reflect impaired adhesion, cytoskeletal contraction, or stress-induced loss of spreading. (D’Urso et al., 2024).

At Day 3, *MP*-treated cells exhibited reductions in several spreading-related parameters, indicating progressive contraction of the cell body. In contrast, ZnO*MP*-treated cells maintained dimensions closer to untreated controls. Preservation of cell area and perimeter suggests maintenance of focal adhesion maturation and intracellular actomyosin tension, both of which are essential regulators of MSC fate determination (Wu et al., 2022). The ability of ZnO*MP* to preserve these descriptors therefore indicates maintenance of a favorable mechanical microenvironment despite prolonged nanoparticle exposure.

Shape-related descriptors provided additional evidence regarding cytoskeletal integrity. Form factor, solidity, and compactness remained relatively stable despite changes in cell size. This observation indicates that treatments did not induce complete cell rounding or catastrophic structural collapse. Instead, MP exposure generated a contracted phenotype while preserving overall cellular geometry. Reduced compactness observed in MP-treated cells is consistent with altered actin organization and force transmission across the cytoskeleton. The recovery of compactness-related descriptors in ZnOMP-treated cultures suggests preservation of cytoskeletal architecture and intracellular mechanical equilibrium (Momotyuk et al., 2025).

Eccentricity and orientation further demonstrated that MSC polarity remained largely preserved across experimental groups. Eccentricity reflects cellular elongation whereas orientation describes directional alignment. The stability of these parameters indicates that treatments affected the extent of cell spreading rather than the fundamental polarity of the cells. Similar observations have been reported during adaptive cellular responses to mechanical and biochemical stress, where cells reduce spreading while maintaining their intrinsic organizational axis (Raman et al., 2022; Bouzid et al., 2019).

The nucleus exhibited even greater sensitivity than the cytoplasm. Nuclear morphology represents a highly responsive indicator of cellular state because the nucleus is mechanically coupled to the cytoskeleton through the linker of nucleoskeleton and cytoskeleton (LINC) complex. Mechanical forces generated within the actin cytoskeleton are transmitted directly to the nuclear envelope, influencing chromatin organization, gene expression, and lineage commitment (Bouzid et al., 2019; Rajendran et al., 2023).

Importantly, nuclear solidity, compactness, and eccentricity remained largely unchanged. Nuclear fragmentation and irregularity are hallmarks of apoptosis and irreversible cellular injury. In contrast, the preservation of nuclear geometry despite reductions in nuclear size indicates an adaptive stress response rather than overt nuclear damage. Morphometric profiling therefore revealed that MP exposure induced significant nuclear plasticity characterized by nuclear compaction while maintaining overall nuclear architecture. Such responses have been associated with chromatin remodeling, altered transcriptional activity, and mechanotransductive adaptation rather than cell death (Bouzid et al., 2019; Raman et al., 2022).

A consistent observation throughout both cellular and nuclear analyses was the protective effect of ZnO*MP*. For multiple descriptors, ZnO*MP*-treated groups exhibited values significantly closer to controls than to MP-treated cultures. Zinc plays fundamental roles in cellular signaling, antioxidant defense, cytoskeletal regulation, and stem cell differentiation (Li et al., 2022; Chen et al., 2024; Peters et al., 2024). The preservation of cell spreading, nuclear dimensions, and morphometric homeostasis observed following ZnO*MP* treatment may therefore reflect improved cellular resilience and maintenance of mechanotransductive signaling pathways.

The functional significance of these observations was further evaluated through osteogenic differentiation assays (Figure 8). Osteogenic cultures exhibited robust mineral deposition after 21 days, confirming successful differentiation. Interestingly, ZnO*MP* promoted mineralization even under basal conditions, suggesting an intrinsic osteoinductive influence. Although ZnO*MP* did not significantly enhance mineralization beyond conventional osteogenic medium, it did not impair extracellular matrix mineralization or calcium deposition. This finding is particularly important because osteogenic competence is a prerequisite for biomaterials intended for musculoskeletal regeneration. Zinc is known to regulate multiple pathways involved in osteoblast differentiation and bone formation, including RUNX2 signaling, alkaline phosphatase activity, and extracellular matrix maturation (Li et al., 2022; Zhu et al., 2024). The preservation of osteogenic differentiation observed in the present study therefore reinforces the biocompatibility of ZnOMP and supports its potential utility in regenerative medicine.

**Figure 8:**
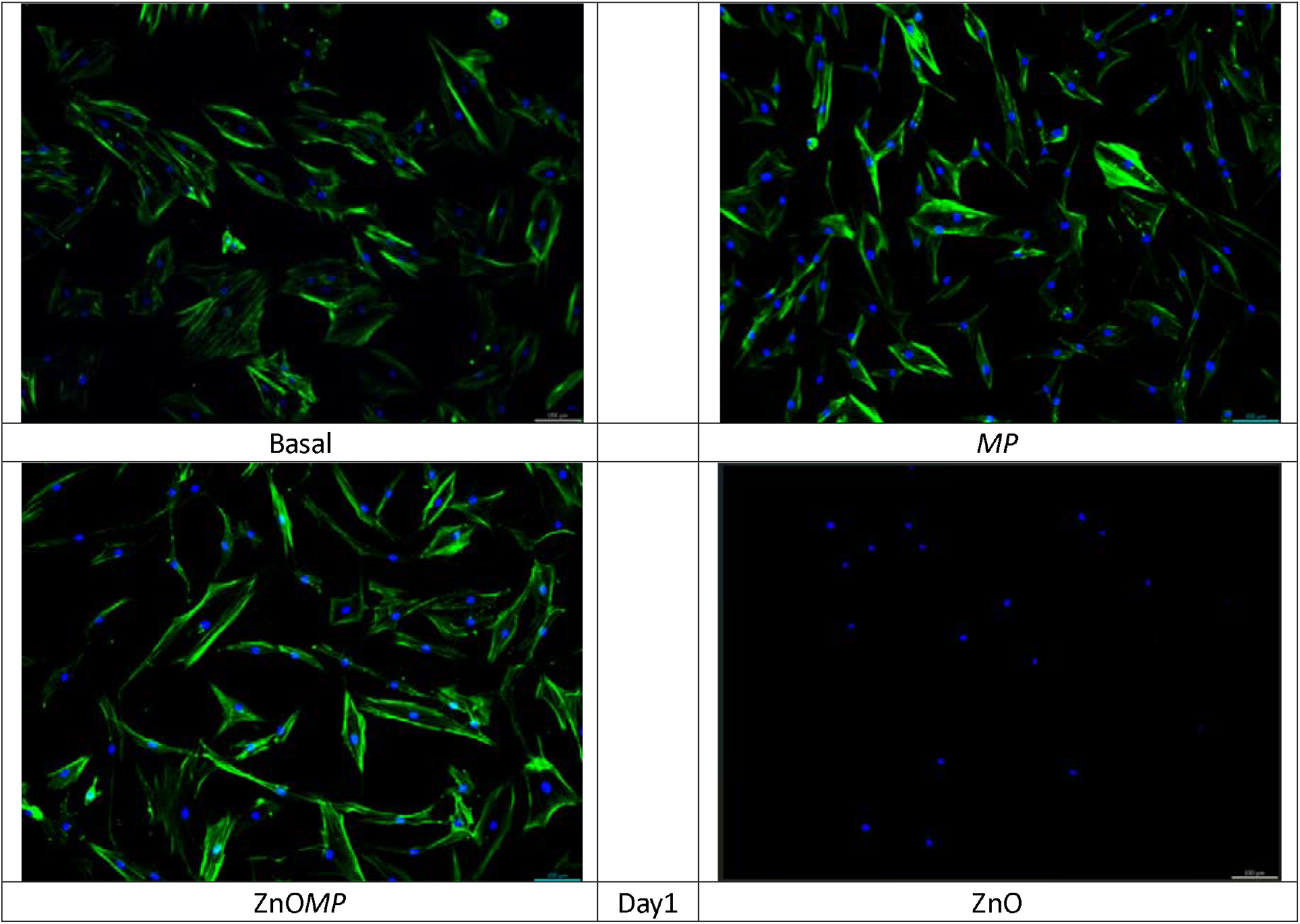
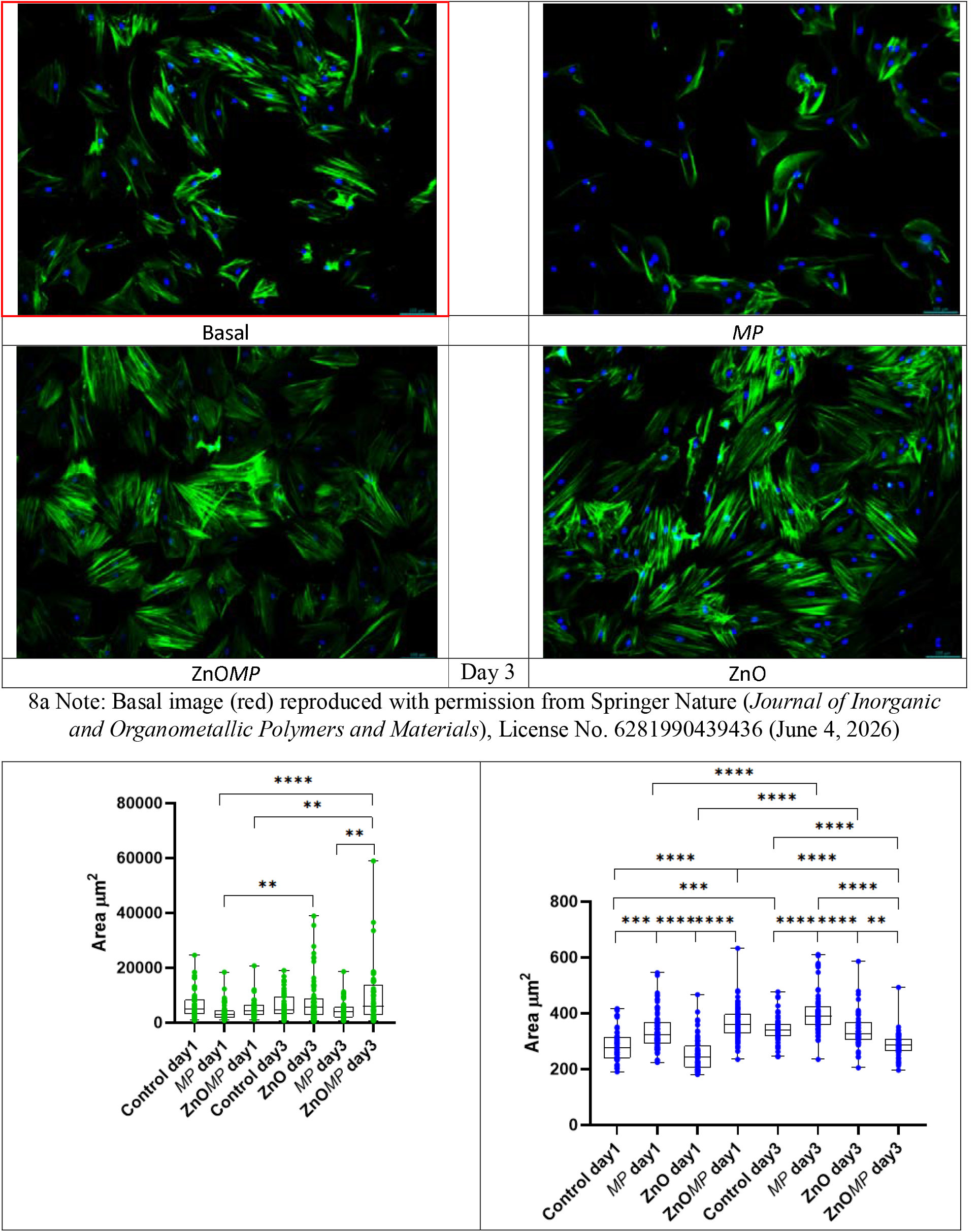
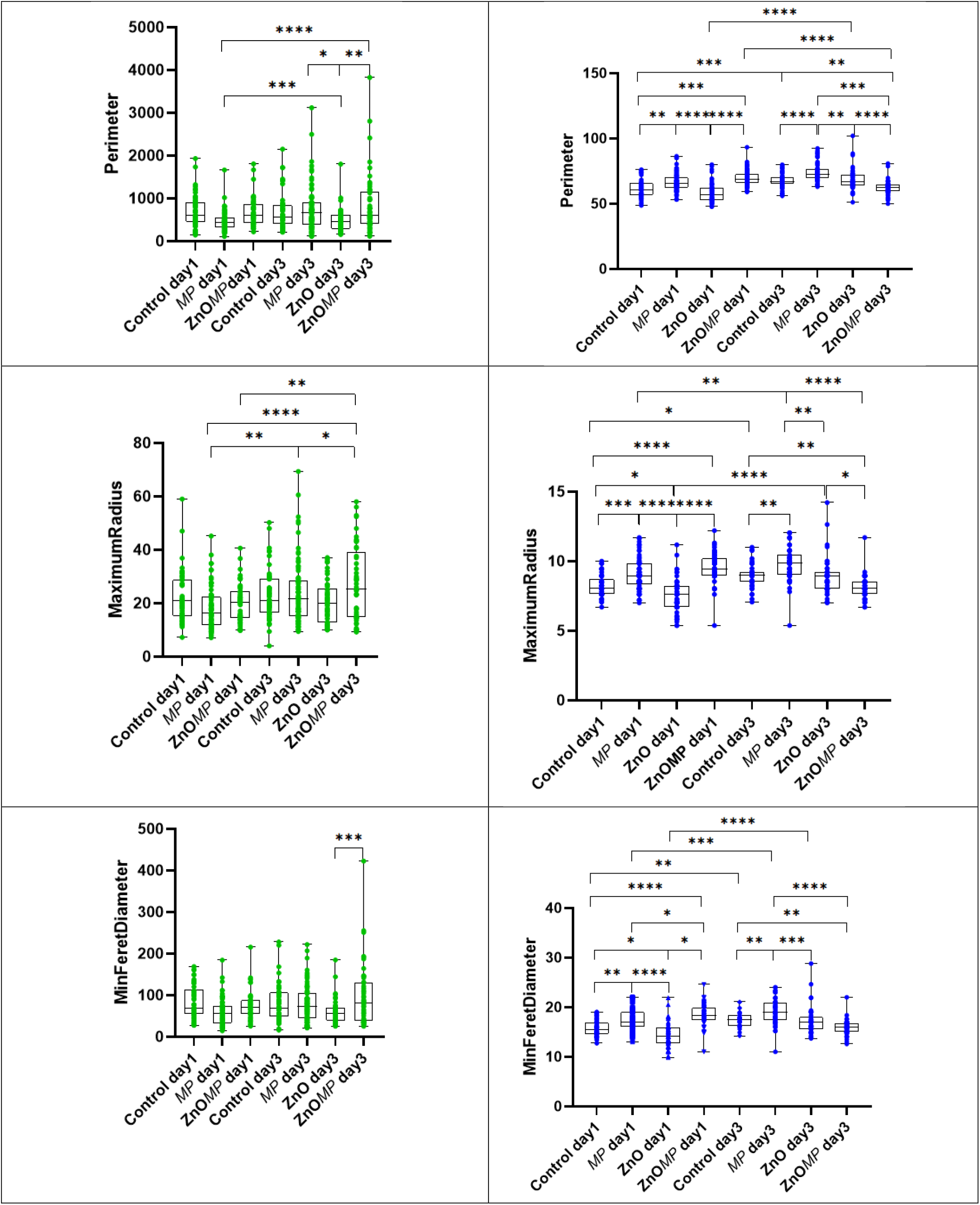
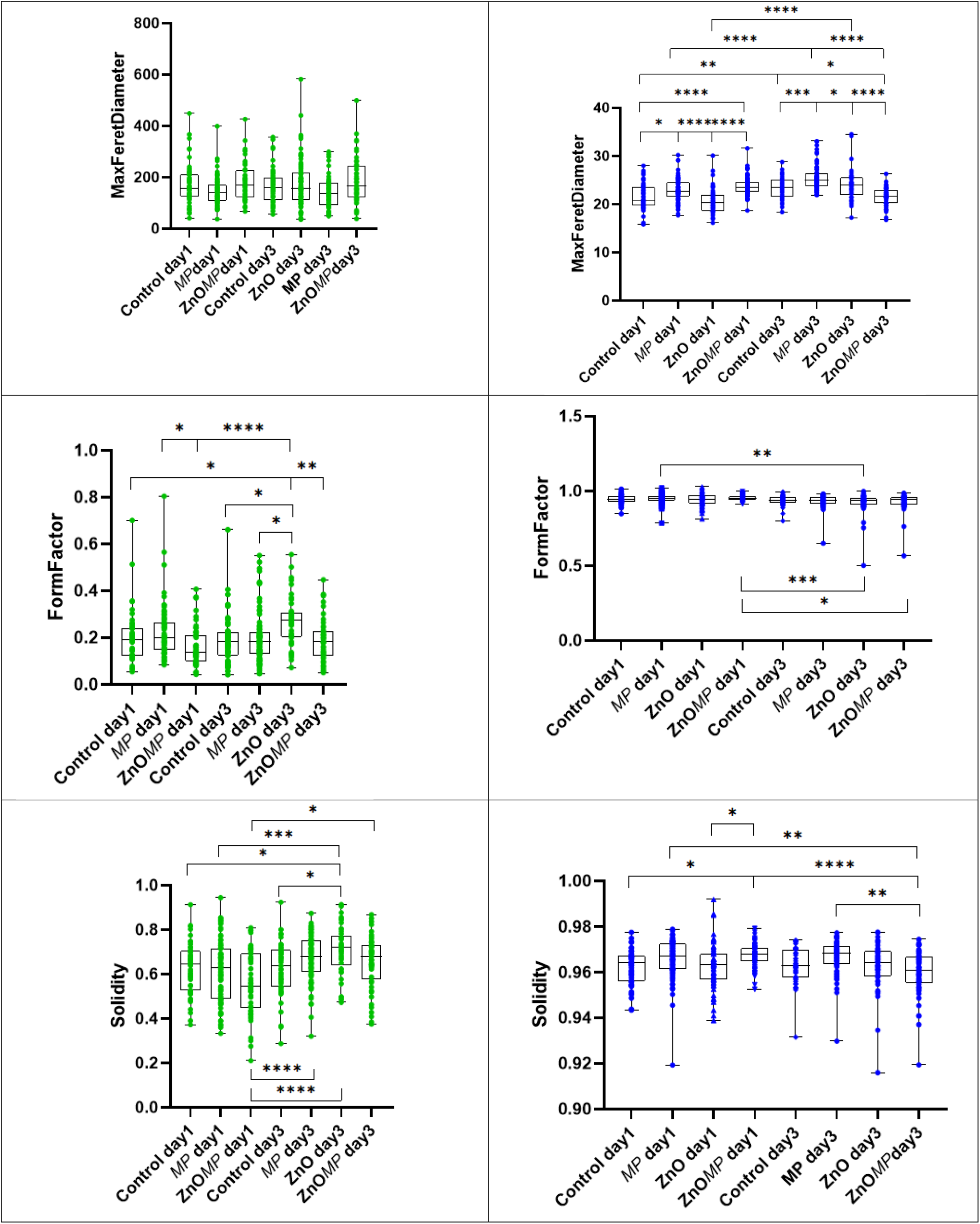
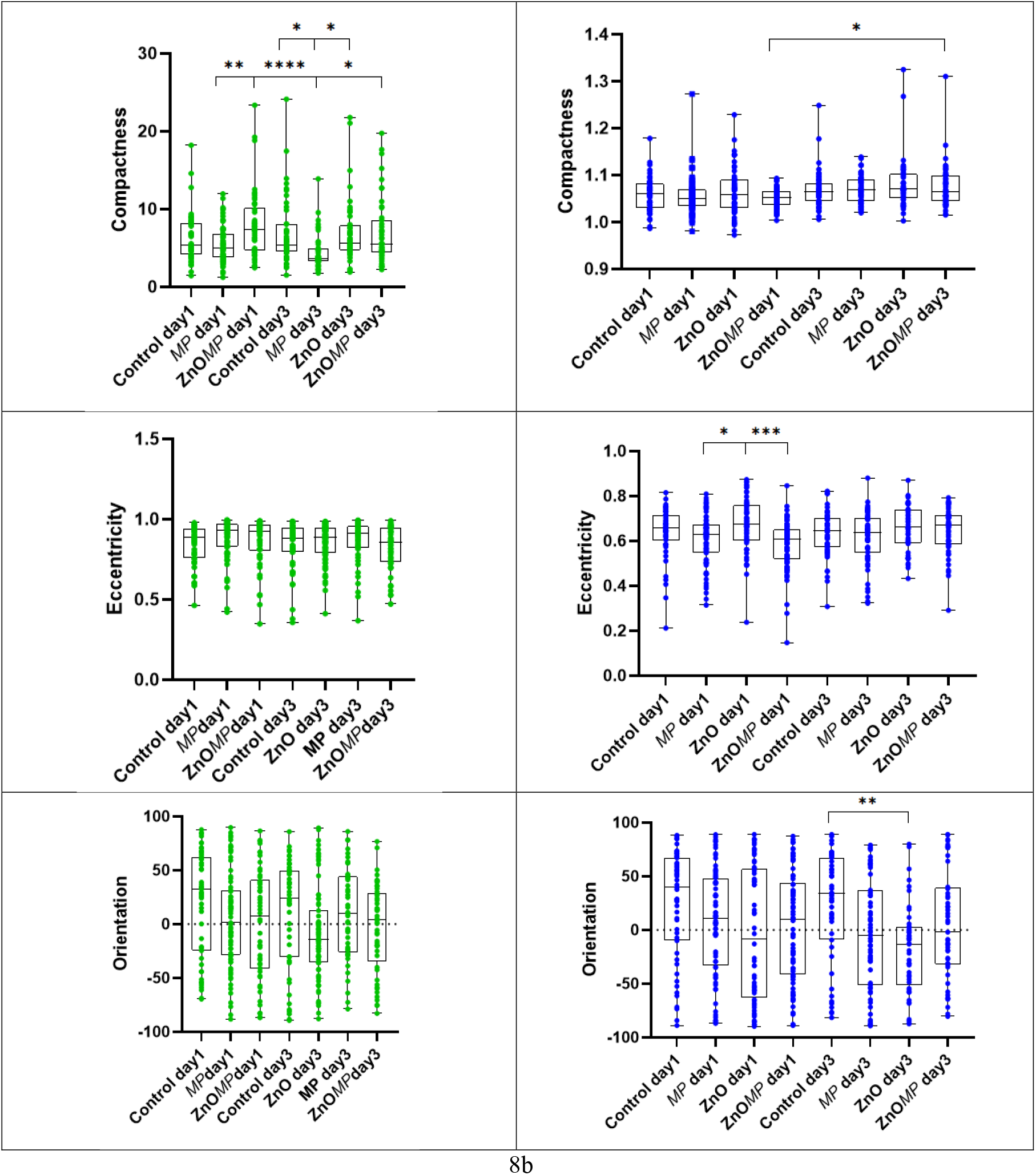
8a Representative composite images of cells stained with Phalloidin-iFluor 488 conjugate to stain actin filaments within the cytoplasm (green) and DAPI to stain the nucleus (blue), scale bar 100 μm at day1 and day3. 8b analysis of single-cell morphological descriptors (area, perimeter, maximumradius maxferetdiameter, minferetdiameter, compactness, solidity, eccentricity, formfactor, orientation) exported from CellProfiler pipeline at day1 and day3. Two way ANOVA was used for comparisons with * = P<0.05, ** = P<0.01, *** = P<0.001, **** = P<0.0001.

**Figure 9:**
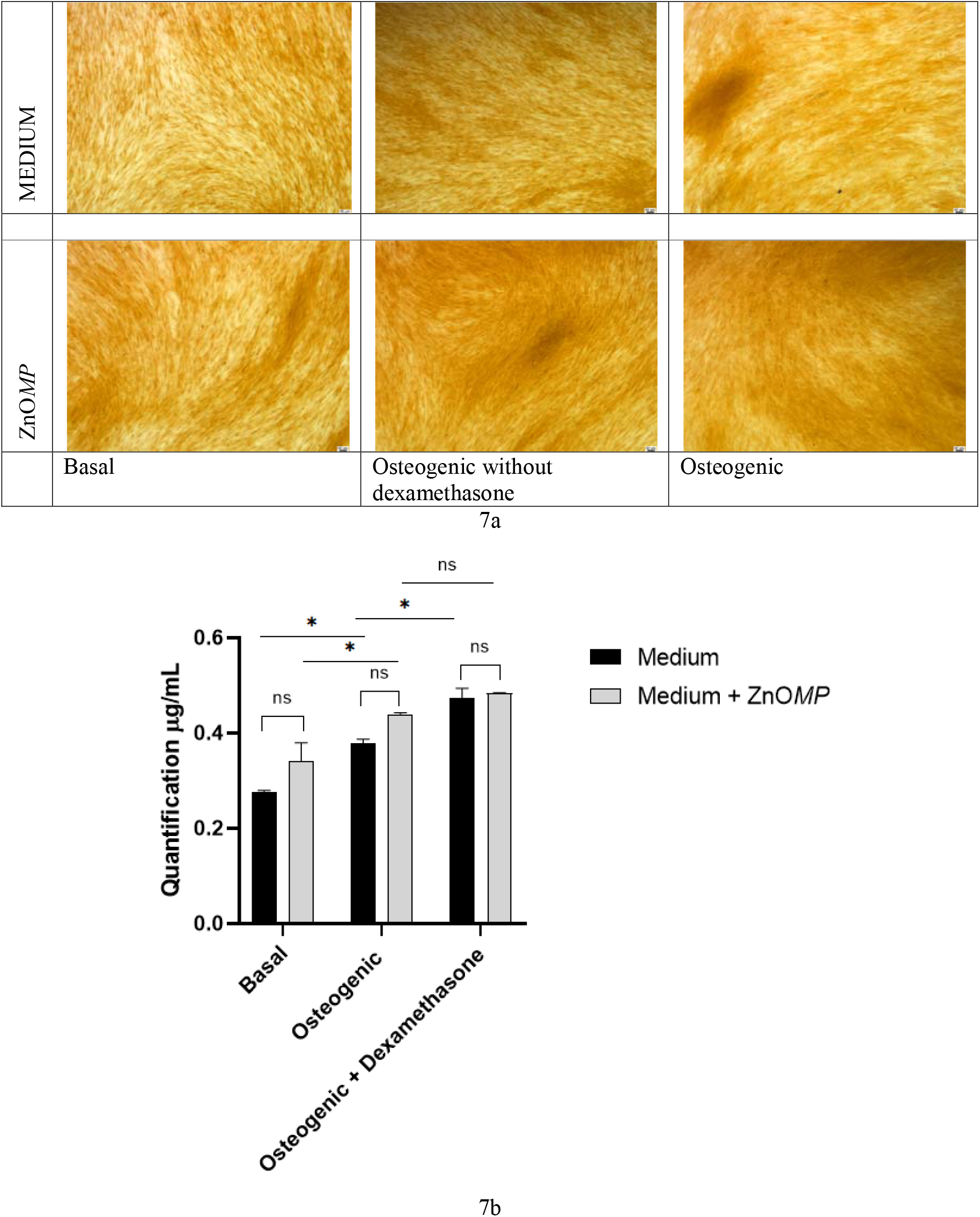
9a Representative images of Alizarin Red S staining at day 21 of basal, and osteogenic culture. Scale bar: 100 μm. 9b Osteogenic potential assessed by quantification of incorporated Alizarin Red S dye at day 21. Cells used from one donor, standard error of the mean and two-way ANOVA is shown

Collectively, these findings demonstrate that *Mimosa pudica* extract induced morphometric and structural alterations but also severe cytotoxicity when MTT experiment is considered. High-content morphological profiling proved particularly sensitive for detecting these early responses, revealing substantial modifications in cellular and nuclear architecture. In contrast, ZnO*MP* preserved cellular morphology, maintained mechanotransductive homeostasis, supported metabolic activity, and retained osteogenic competence. The convergence of physicochemical characterization, viability measurements, morphometric analyses, and differentiation assays identifies ZnO*MP* as a promising phytogenic nanomaterial for musculoskeletal tissue engineering and regenerative medicine.

## Conclusion

This study demonstrates that *Mimosa pudica* leaf extract can serve as an effective biogenic platform for the synthesis of zinc oxide nanoparticles with physicochemical characteristics suitable for biomedical applications. FTIR, PXRD, SEM, EDS, and TEM analyses confirmed the formation of phytochemical-capped ZnO nanoparticles, indicating that plant-derived metabolites actively participated in nanoparticle reduction and stabilization.

Biological evaluation showed that ZnO*MP* were well tolerated by human bone marrow mesenchymal stromal cells and supported higher cellular metabolic activity than either the plant extract alone or chemically synthesized ZnO nanoparticles. Beyond conventional viability measurements, high-content morphometric analysis revealed that exposure to *Mimosa pudica* extract altered shape organization, and nuclear dimensions, effects consistent with cytoskeletal stress. In contrast, ZnO*MP* attenuated many of these alterations and maintained cellular and nuclear characteristics closer to those of untreated controls, suggesting improved cellular adaptation and preservation of structural integrity.

Importantly, ZnO*MP* did not compromise osteogenic differentiation. Mesenchymal stromal cells retained their capacity to deposit a mineralized extracellular matrix under osteogenic conditions, demonstrating preservation of functional competence despite prolonged nanoparticle exposure. Although ZnO*MP* did not significantly enhance mineralization, the maintenance of osteogenic potential represents a critical requirement for regenerative applications materials. Collectively, these findings identify *Mimosa pudica*-derived ZnO nanoparticles as biocompatible nanomaterials capable of supporting mesenchymal stem cell function while preserving osteogenic capacity. Their favorable biological profile warrants further investigation in *in vitro* models and preclinical studies targeting musculoskeletal tissue regeneration.

## Author Contributions

Eya’ane Meva F, Janiak C; Writing – review & editing, Conceptualization, Supervision, Investigation, Funding acquisition. Djuidje AG, Belle Ebanda Kedi P, Ntoumba AA, Marcus N. A. Fetzer; Writing – review & editing, Conceptualization, Visualization, Intrumentation. Belle Ebanda Kedi P, Fonye Nuyfoni G, Chimi Tchoutchang G, Nanga CC, Mintang Fongang UA, Tako Djimefo AK, Tabearuh Ayuk BT, Evouna MID; Writing– review & editing, Investigation, Data curation, Animal models, Instrumental support.

## Limitations

Several limitations of the present study should be acknowledged. First, the biological investigations were conducted using BM-MSCs isolated from a limited number of human donors (2). Second, the morphometric analyses were performed on a relatively limited number of cells per experimental condition (approximately 50–80 cells). Finally, the present work was restricted to in vitro evaluation of BM-MSC viability, morphology, and osteogenic differentiation. Although the findings demonstrate favorable cytocompatibility and preservation of osteogenic competence, additional studies are required to investigate the long-term biological behavior, biodistribution, immunomodulatory effects, and regenerative performance of ZnO*MP* in physiologically relevant three-dimensional models and in vivo musculoskeletal regeneration systems.

## Ethical approval

The study was conducted according to the guidelines of the Declaration of Helsinki and approved by the Ethics Committee of the University of Würzburg (187/18). Informed consent was obtained from all subjects involved in the study.

## Acknowledgements

Authors thank Prof Marietta Herrmann, Musculoskeletal Cell Biology Group, Institute for Functional Materials and Biofabrication, University Hospital Würzburg and Chair for Orthopedics, Julius-Maximilians-Universität Würzburg, Röntgenring 11, 97070 Würzburg, Germany, for the mesenchymal stem cells experiments. Support of the Imaging Core Facility, Biocenter, University of Würzburg, Germany for the JEOL JEM-2100 Transmission Electron Microscope funded by the Deutsche Forschungsgemeinschaft (DFG, German Research Foundation) - 218894163 is acknowledged.

## Data Availability

Data will be provided by the authors upon reasonable request.

---

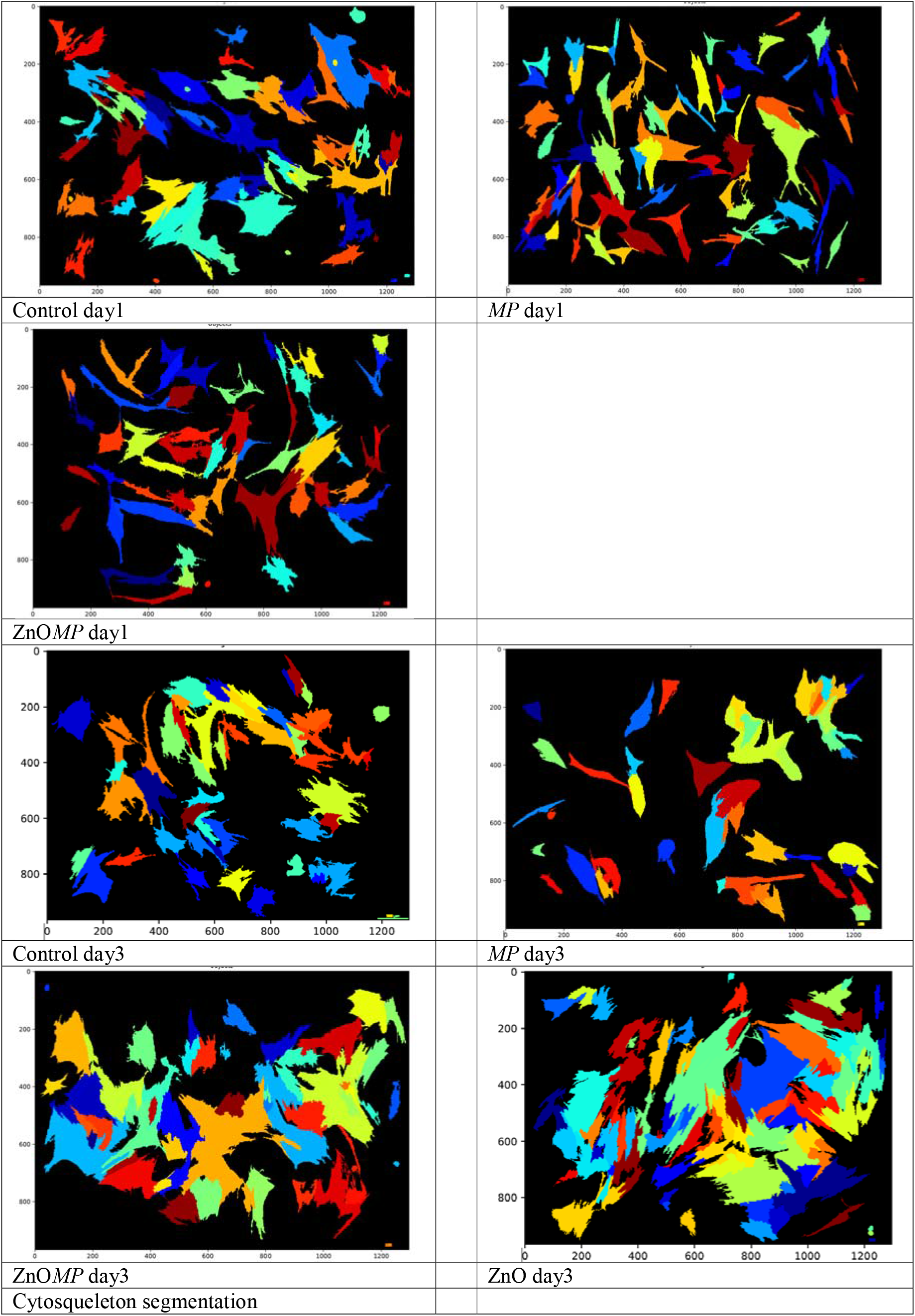

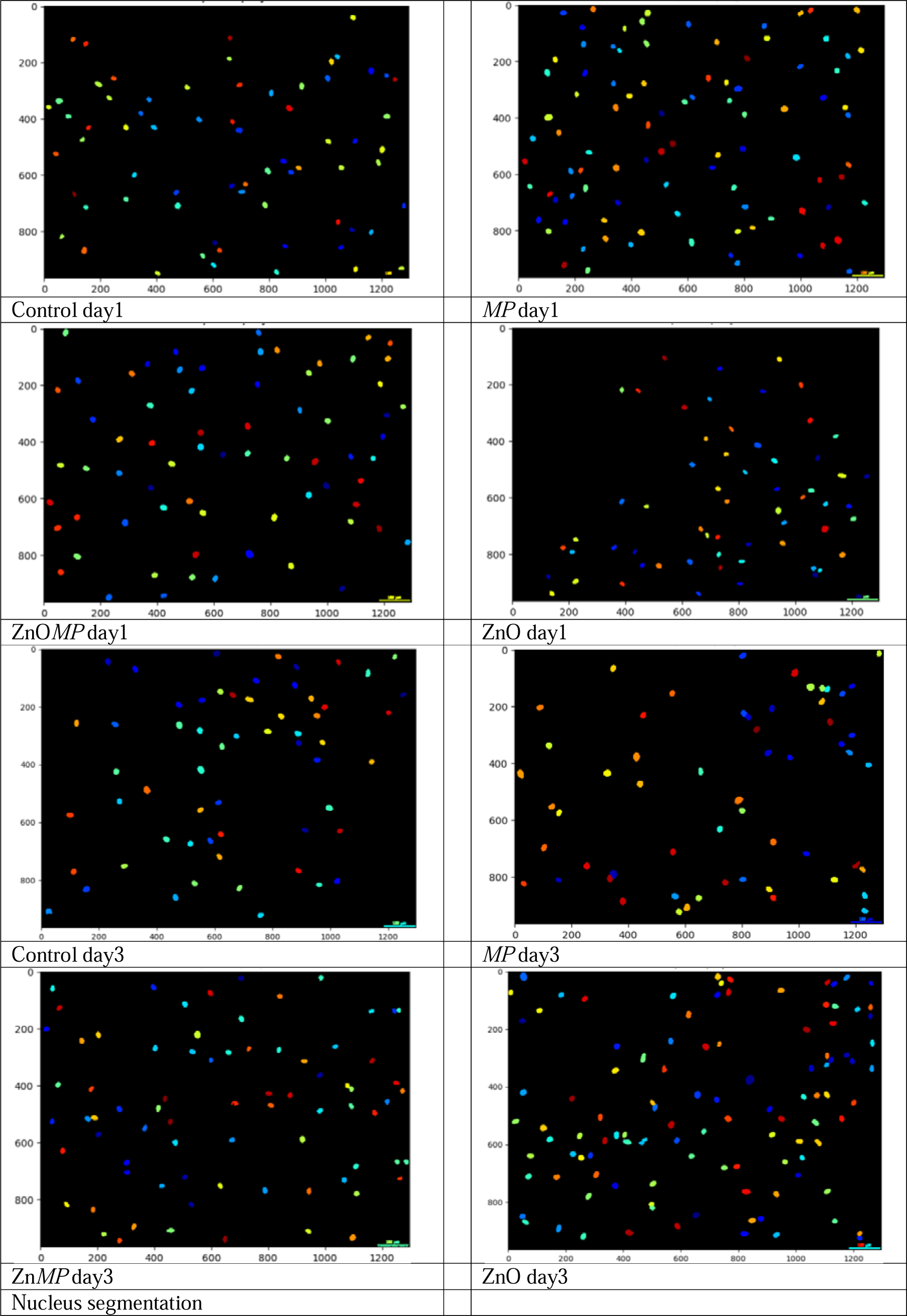

